# STREAMLINING HISTOPATHOLOGY: INSTAPRESERVE TISSUESAFE A SINGLE-STEP FIXATIVE THAT IMPROVES TISSUE QUALITY AND WORKFLOW EFFICIENCY

**DOI:** 10.1101/2025.05.01.651707

**Authors:** Rajani Kanth Vangala, Anand Babu Vangala, Saranya Kanukollu, M S Pratibha, S Avinash, Elango E Murugaian

**Affiliations:** Neuome Technologies Private Limited, Electronic City, Bangalore, India

**Author notes:** Communication author. Communication address: Dr. Elango Murugaian, Neuome Technologies Private Limited, V V Tower, Beratene Agrahara, Begur Hobli, Hosur Road, Electronic City, Bengaluru - 560100 Karnataka.

## Abstract

In clinical pathology, tissue processing is crucial in generating a valuable report for faster diagnosis. Formalin-fixed paraffin-embedded tissues are the standard specimens for clinical and molecular analyses. Though the demand persists for faster diagnosis, neutral buffered formalin (NBF) -based fixation and processing have not been replaced with a simple, faster, and user-friendly method. The alcohol-based fixatives were evaluated as an alternative to formalin, yet none have replaced formalin for various reasons. InstaPRESERVE Tissuesafe is an alcohol-based, biodegradable, and user-friendly fixative solution. The clinical specimens are well preserved and suitable for histopathology, genomics, and proteomics studies. The efficacy of instaPRESERVE Tissuesafe with ethanol or isopropyl alcohol and a modified tissue processing method without alcohol dehydration steps was studied.

The pig’s organs, brain, and liver tissues were fixed with NBF and InstaPRESERVE Tissuesafe solutions (ethanol/isopropyl alcohol). Based on the alcohol content of instaPRESERVE Tissuesafe, the dehydration step(s) were skipped during tissue processing. Serial sections from fixed tissues were processed and stained with H&E. The intactness of the sections with instaPRESERVE Tissuesafe fixation was comparable to that of NBF fixation. No autolysis/ decomposition was observed with InstaPRESERVE Tissuesafe (both) and NBF. The H&E staining scoring was satisfactory with well-preserved cell boundaries and organelles in both liver and brain tissue sections.

The key factors of any fixative are its penetration, temperature, and time taken for fixation. In this study, the penetration rate of InstaPRESERVE Tissuesafe with either ethanol or isopropyl alcohol was the same as NBF within 6 to 12 hrs. The results indicated that increased concentration of alcohol in instaPRESERVE Tissuesafe did not affect the histology, and overall quality of the tissue sections processed was comparable to formalin fixation. The cytoplasmic and nuclear details, erythrocyte integrity, and lymphocytic appearance were well maintained. The time was reduced to 6 hrs from 9 hrs of the standard method.

This study has addressed the modification of tissue processing with reduced time in histopathology with instaPRESERVE Tissuesafe. The outcome of this study has shown that instaPRESERVE Tissuesafe may be an alternative to formalin and helps in faster diagnosis.

## Introduction

In clinical pathology, rapid diagnosis is increasingly desirable to meet the needs of clinicians. The histology techniques are time-consuming, resulting in delayed diagnosis. The processing of biospecimens from the point of tissue removal to embedding is laborious. After archival, the tissue undergoes preparation of sections, usually fixation with a suitable supporting medium. The tissue processing stages are fixation, dehydration, clearing, impregnation, and embedding for designated time durations [1]. The current methods are accepted globally with reproducibility and relatively low expense. Though the demand persists for faster or early reporting, neutral buffered formalin-based fixation and processing have not been replaced with a simple, faster, and user-friendly method [2]. Immunohistochemistry and molecular methods are validated on formalin-fixed material; hence difficult to introduce a new fixative into histopathological practice. Another significant issue with formalin is RNA extraction [3]. RNA extracted from FFPE tissue often experiences strand breakage and crosslinking, leading to inconsistent quality and yield.

In pathology laboratories, large amounts of formalin are used, and technicians and pathologists are exposed to a dilute solution of hazardous formaldehyde, often undervaluing its side effects [4, 5]. The existing tissue processing is reliable and cost-effective, but it is associated with disadvantages like reagent toxicity and extended processing duration, thus delaying diagnosis. The rapid manual tissue processing method is allied with harmful chemicals and higher tissue distortion and shrinkage. This emphasizes the need for newer short-processing techniques. Several other fixatives have been tested as non-hazardous and nucleic acid-preserving alternatives, but none have displaced formalin as the standard fixative for various reasons [6-9]. The demand for faster or early reporting is increasing, and newer techniques such as rapid manual and microwave processing are being validated. Each of them is unique with its own set of advantages and disadvantages.

Swine is a suitable animal model used in translational research, preclinical toxicologic testing of pharmaceuticals, and surgical training [10, 11]]. Due to the similarity to humans in anatomy, genetics, and physiology, pigs are potentially better than other large animal species. Humans and swine are omnivorous, and their organs generally share common functional and metabolic features. Hence, brain and liver tissues from a pig were used in this study.

InstaPRESERVE Tissuesafe is an alcohol-based, biodegradable, and user-friendly fixative solution to preserve biospecimens at room temperature. The clinical specimens are well preserved and suitable for histopathology, genomics, and proteomics studies [12]. Since instaPRESERVE Tissuesafe is an alcohol-based solution, ethanol and isopropyl alcohol were evaluated for their efficacy in replacing formalin in histopathology. A modified tissue processing method without ethanol dehydration steps was scrutinized to enhance faster diagnosis with reduced processing time.

## Materials and Methods

### Organ specimen collection and fixation

InstaPRESERVE Tissuesafe was developed with ethanol and validated [12]. In this study, modified instaPRESERVE Tissuesafe, wherein ethanol was replaced with isopropyl alcohol, was included. Additionally, formulations with increasing percentages of ethanol and isopropyl alcohol from 70%, 80%, and 90% were prepared (Table 1). The standard 10% neutral buffered formalin was used as a control.

**Table 1:** Fixative solutions and formulations.

| S.No | Fixative solution | Ethanol % | Isopropyl alcohol % |
| --- | --- | --- | --- |
| 1 | IP-E70 | 70 | - |
| 2 | IP-E80 | 80 | - |
| 3 | IP-E95 | 95 | - |
| 4 | IP-I70 | - | 70 |
| 5 | IP-I80 | - | 80 |
| 6 | IP-I95 | - | 95 |

The pig’s organs, brain, and liver were collected freshly from a local slaughterhouse. After removing blood stain with water, the organs were cut into 4 cubic centimeter pieces. The pieces representing the same anatomical regions were fixed with buffered formalin and InstaPRESERVE Tissuesafe solutions (1:7 w/v ratio). The specimens were sent to the histopathology lab within 1 hour.

### Paraffin embedding and microtomy

After 12 hours of fixation, serial sections were made, grossed, and processed in an automated tissue processing System (Leica Biosystems, Germany). Based on the alcohol content of instaPRESERVE Tissuesafe, the dehydration step was skipped during dehydration (Table 2). For example, while processing the tissue fixed with instaPRESERVE Tissuesafe with 70% ethanol (IP-E70), the first step, 70% ethanol, was omitted. For instaPRESERVE Tissuesafe with 80% ethanol, the first two steps, 70% and 80%, were omitted. In control, NBF fixed tissue sections were processed like instaPRESERVE Tissuesafe with ethanol. For the tissues fixed with instaPRESERVE Tissuesafe containing isopropyl alcohol, ethanol was replaced with isopropyl alcohol. The standard dehydration procedure was followed in the control experiment of the three fixative solutions.

**Table 2:** Modification of dehydration in tissue processing.

| S.No | Samples | Dehydration steps |  |  |  |
| --- | --- | --- | --- | --- | --- |
|  |  | 70% | 80% | 95% | 100% |
|  | **Ethanol** |  |  |  |  |
| 1 | IP-E70 | - | Yes | Yes | Yes |
| 2 | IP-E80 | - | - | Yes | Yes |
| 3 | IP-E95 | - | - | - | Yes |
| 4 | IP-E control | Yes | Yes | Yes | Yes |
| 5 | 4% NBF-E70% | - | Yes | Yes | Yes |
| 6 | 4% NBF-E80 | - | - | Yes | Yes |
| 7 | 4% NBF-E95 | - | - | - | Yes |
| 8 | 4% NBF-control | Yes | Yes | Yes | Yes |
|  | **Isopropyl alcohol** |  |  |  |  |
| 9 | IP-I70 | - | Yes | Yes | Yes |
| 10 | IP-I80 | - | - | Yes | Yes |
| 11 | IP-I95 | - | - | - | Yes |
| 12 | IP-I control | Yes | Yes | Yes | Yes |

**Table 3:** H&E staining scoring of the brain and liver tissue sections fixed with NBF and instaPRESERVE Tissuesafe (ethanol and isopropyl alcohol).

| Fixative Solution | Liver | Brain |
| --- | --- | --- |
| IP-E70 | 3+3+3=9 | 3+3+3=9 |
| IP-E80 | 3+3+3=9 | 3+3+3=9 |
| IP-E95 | 3+3+3=9 | 3+3+3=9 |
| IP-E control | 3+3+3=9 | 3+3+3=9 |
| 4% NBF-E70% | 3+3+3=9 | 2+3+3=8 |
| 4% NBF-E80 | 3+3+3=9 | 3+3+3=9 |
| 4% NBF-E95 | 3+3+3=9 | 3+3+3=9 |
| 4% NBF-control | 3+3+3=9 | 3+3+3=9 |
| IP-I70 | 3+3+3=9 | 3+3+3=9 |
| IP-I80 | 3+3+3=9 | 3+3+3=9 |
| IP-I95 | 3+3+2=8 | 1+3+3=7 |
| IP-I control | 3+3+3=9 | 1+3+3=7 |

Following dehydration, the tissue sections were cleared with xylene and embedded in paraffin blocks. From each block, 1-1.5 µm sections were prepared using a sliding microtome and transferred to slides.

### H&E staining and scoring

The slides were subjected to routine hematoxylin and eosin (H&E) stain protocol. The microscopic scoring was based on 1. Intactness of sections 2. Nuclear staining 3. Cytoplasmic staining. Under each category, the score was Good – 3, Moderate – 2, Poor - 1, with an overall score of 8 or 9 – Satisfactory, and 7 or less – Unsatisfactory. The intactness scoring was based on membrane damage, cell proposition, absence of organelles, and artifacts due to staining, shrinkage, or sectioning.

## Results

The tissue specimens fixed with InstaPRESERVE Tissuesafe (ethanol or isopropyl alcohol) were in good condition without discoloration and degradation from collection until the study completion. The formalin-fixed tissues exhibited regular discoloration. The penetration time of formalin and InstaPRESERVE Tissuesafe (both) was around 1mm per hour. The scoring was satisfactory for all sections except for 95% isopropyl alcohol (IP-I95) and its control (Table 2). No autolysis/ decomposition was observed from the tissues fixed with InstaPRESERVE Tissuesafe (both) and NBF. The intactness of the tissue sections with InstaPRESERVE Tissuesafe (both) fixation was comparable to that of NBF-fixed tissues. The boundaries were well maintained, and none of the sections showed any membrane damage, shrinkage, or less staining except for the brain tissue fixed IP-I95 and the control. Though the nuclear and cytoplasmic staining was good, the intactness of the sections was not satisfactory.

The microscopic analysis of the H&E-stained slides showed well-preserved cell boundaries and organelles in both liver and brain tissue sections. In the liver, the RBCs are well appreciated with InstaPRESERVE Tissuesafe fixation with either ethanol or isopropyl alcohol (Fig. 1). The liver lobule exhibited spokes of wheel arrangement of a central vein surrounded by hepatocytes. The lobule was presented with an edematous boundary, separated from the surrounding parenchyma. While the intactness of brain tissue sections was satisfactory with 70% and 80% isopropyl alcohol containing instaPRESERVE Tissuesafe, unsatisfactory intactness was observed with 95% and the control sections.

**Figure 1:**
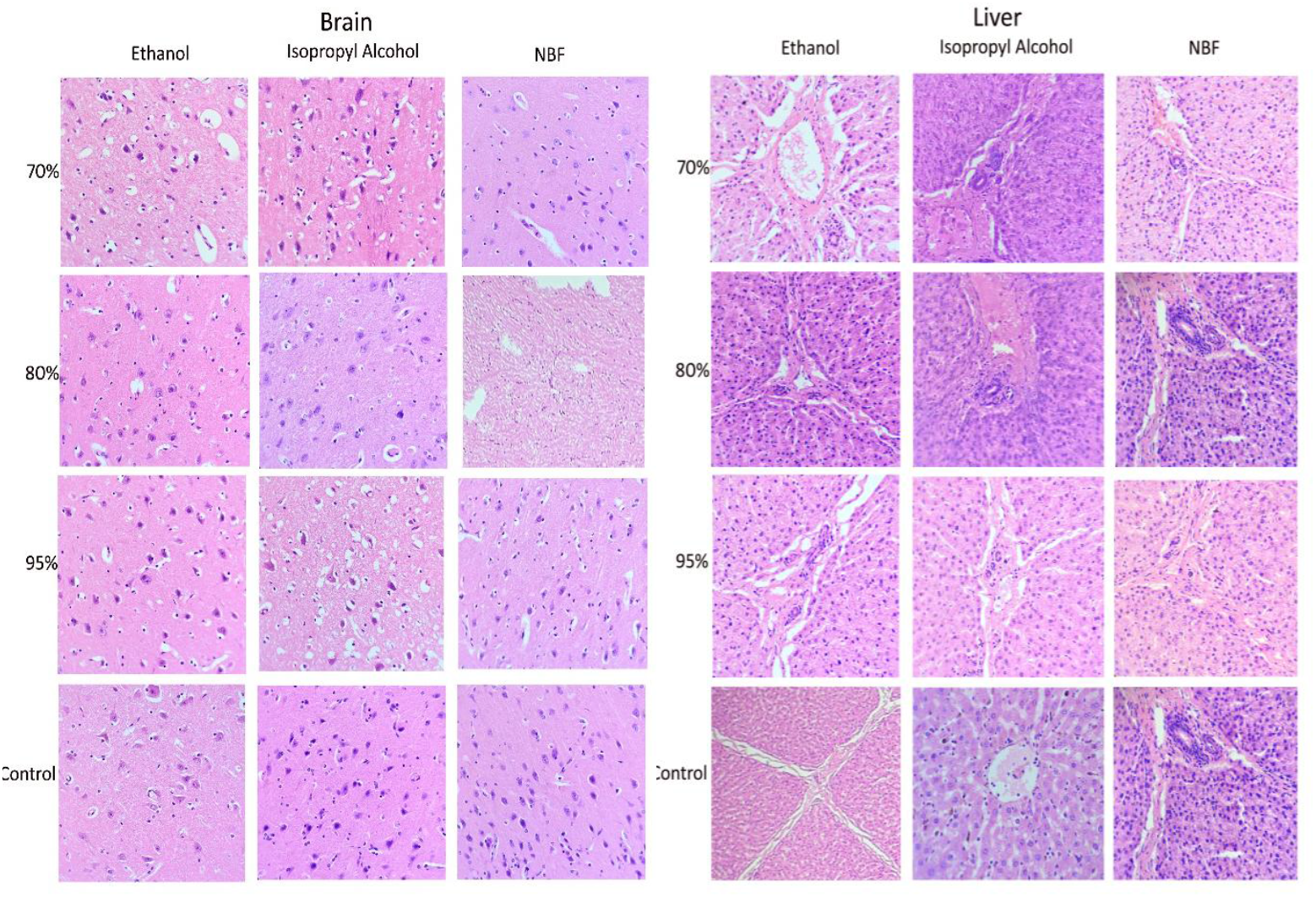
H&E staining analysis of InstaPRESERVE Tissuesafe (ethanol or isopropyl alcohol) and formalin-fixed liver and brain.

## Discussion

The crucial steps in histopathology are the selection of a fixative and the tissue preparation. Formalin-fixed paraffin-embedded tissues are the standard specimens for clinical and molecular analyses. Due to the cross-linking nature of formalin, the extraction efficiency is reduced with poor quality of DNA, RNA, and proteins [3, 13, 14]. Commercially available molecular fixatives have been developed to obtain preservation of morphology and staining results similar to formalin [15]. While formaldehyde has fixation efficacy in NBF, its hydrated form, methylene glycol, has high penetration efficiency. The alcohol-based fixatives evaluated as an alternative to formalin caused protein coagulation and tissue contraction, due to dehydrating the proteins. To prevent coagulation, many of the alcohol-based fixatives are added with protein stabilizers or chloroform and acetic acid. Though not limiting the diagnosis, Paxgene-fixed tissue was reported to have increased eosinophilia [16]. Hemolysis and shrinkage of tissues are associated with many molecular fixatives like RCL2, PAXgene, FineFix, and F-Solv [17]. InstaPRESERVE Tissuesafe, as an alcohol-based fixative solution, contains stabilizers to protect the tissue anatomy, morphology, and cellular components [18]. The brain and liver tissues from the pig fixed with InstaPRESERVE Tissuesafe (ethanol or isopropyl alcohol) did not show shrinkage, discoloration, or cell lysis.

The key factors of any fixative are its penetration, temperature, and time taken for fixation. The tissue processing method is critical in generating a valuable report for faster diagnosis [19]. The thumb rule for tissue penetration is 1 mm/h with a recommended fixation time of 24 hrs for NBF-fixed specimens. In this study, the penetration rate of InstaPRESERVE Tissuesafe with either ethanol or isopropyl alcohol was the same as NBF within 6 to 12 hrs. Rapid processing of tissues in histology analysis is more desirable for timely diagnosis and intraoperative consultations. Many methods are validated for reduced tissue fixation, processing, sectioning, and staining [14, 20, 21]. NBF fixed tissues are sequentially dehydrated with ethanol, cleared with xylene, and embedded in paraffin wax. The standard processing time is more than 9 hrs, which largely impacts the delay in diagnosis. InstaPRESERVE Tissuesafe is an alcohol based fixative solution; hence, reduced dehydration time was evaluated in this study. InstaPRESERVE Tissuesafe with increasing concentrations (70% to 95%) of ethanol or isopropyl alcohol was used as a fixative solution. As per the concentration of alcohol used for fixation, the dehydration steps were omitted. The results indicated that an increased percentage of alcohol in instaPRESERVE Tissuesafe did not affect the histology, and overall quality of the tissue sections processed was comparable to formalin fixation. The cytoplasmic and nuclear details, erythrocyte integrity, and lymphocytic appearance were well maintained. The time was reduced to 6 hrs from 9 hrs of the standard method.

## Conclusion

This study has addressed the modification of tissue processing in histopathology using instaPRESERVE Tissuesafe with either ethanol or isopropyl alcohol. This alcohol based fixative solution has been shown to provide comparable results with formalin fixation with reduced time (9 hrs to 6 hrs). Since both ethanol and isopropyl alcohol in instaPRESERVE Tissuesafe have exhibited similar results, this alcohol-based fixative solution may be optimized based on the user’s convenience. In principle, this study has shown that instaPRESERVE Tissuesafe may be considered an alternative to formalin and helps in faster diagnosis.

## Contributors

All authors have been part of planning, conducting, and reporting this work.

